# Hypoxia-sensing by the Histone Demethylase UTX (*KDM6A*) Controls Colitogenic CD4^+^ T cell Fate and Mucosal Inflammation

**DOI:** 10.1101/2023.07.27.550746

**Authors:** Mandy I. Cheng, Lee Hong, Bryan Chen, Scott Chin, Christopher R. Luthers, Christian Bustillos, Shehzad Z. Sheikh, Maureen A. Su

## Abstract

Hypoxia is a feature of inflammatory conditions [e.g., inflammatory bowel disease (IBD)] and can exacerbate tissue damage in these diseases. To counteract hypoxia’s deleterious effects, adaptive responses have evolved which protect against hypoxia-associated tissue injury. To date, much attention has focused on hypoxia-activated HIF (hypoxia-inducible factor) transcription factors in these responses. However, recent work has identified epigenetic regulators that are also oxygen-sensitive, but their role in adaptation to hypoxic inflammation is currently unclear. Here, we show that the oxygen-sensing epigenetic regulator UTX is a critical modulator of colitis severity. Unlike HIF transcription factors that act on gut epithelial cells, UTX functions in colitis through its effects on immune cells. Hypoxia results in decreased CD4^+^ T cell IFN-γ production and increased CD4^+^ regulatory T cells, and these findings are recapitulated by T cell-specific UTX deficiency. Hypoxia impairs the histone demethylase activity of UTX, and loss of UTX function leads to accumulation of repressive H3K27me3 epigenetic marks at IL12/STAT4 pathway genes (*Il12rb2, Tbx21,* and *Ifng*). In a colitis mouse model, T cell-specific UTX deletion ameliorates colonic inflammation, protects against weight loss, and increases survival. Together these findings implicate UTX’s oxygen-sensitive histone demethylase activity in mediating protective, hypoxia-induced pathways in colitis.

## Introduction

Hypoxia occurs when oxygen demand outstrips supply and is a feature of inflammatory bowel disease (IBD)^1,2^ and other inflammatory conditions. The consequences of hypoxia can be potentially dire, with sustained hypoxia in pathologic sites leading to metabolic crisis and cell death. To adapt to these hypoxic threats, oxygen sensing mechanisms have evolved that mediate a protective response to low oxygen conditions. These responses therefore represent potential pathways that may be harnessed for therapeutic benefit in inflammatory diseases. Indeed, pharmacologic approaches that augment these responses have shown promise in multiple mouse models of colitis^3–5^ and are now under active development for IBD^2^.

To date, studies of hypoxia-induced adaptive responses have focused on pathways mediated by hypoxia-inducible factor (HIF) transcription factors, which promote expression of multiple genes important in adaptation to low oxygen levels^2,6^. Consequently, targeted therapies that stabilize HIF are currently being evaluated as treatments for IBD^1^. For example, a recent double-blind, placebo-controlled trial of GB004, a small molecule stabilizer of HIF-1alpha, demonstrated improved disease activity in ulcerative colitis patients^7^. Thus, HIF stabilization is a promising therapeutic strategy for IBD.

Recently, it has become clear that global hypoxia-induced transcriptional changes can be mediated not only by HIF transcription factors, but also by an oxygen-sensitive epigenetic regulators^8,9^. The role of these epigenetic regulators in hypoxia-associated inflammation, however, is unclear. Here, we demonstrate that impairment of the oxygen-sensitive epigenetic regulator UTX (ubiquitously transcribed tetratricopeptide repeat protein on the X chromosome; encoded by the gene *Kdm6a*) is protective in colitis. Unlike HIF transcription factors, which serve a barrier-protective role in mucosal inflammation, UTX mediates pathways important in colitogenic T cell responses. Hypoxia is associated with decreased CD4^+^ T cell production of IFN-γ^10–13^, a pathogenic cytokine in inflammatory colitis^14–16^, and we find that T cell-specific UTX ablation mimics these changes. Additionally, hypoxia-associated increase in immunosuppressive CD4^+^ CD25^+^ FOXP3^+^ regulatory T cells (Tregs)^17^ is also mirrored by T cell-specific UTX deficiency. Both hypoxia and UTX deletion lead to H3K27me3 (trimethylated histone 3 lysine 27) accumulation in stimulated CD4^+^ T cells, suggesting that hypoxia evokes transcriptional changes through impairment of UTX’s H3K27me3 demethylase activity. Concomitant RNA sequencing and H3K27me3 CUT&Tag demonstrate coordinate UTX-regulated transcription and removal of repressive H3K27me3 marks at IL12/STAT4 pathway genes (*Il12rb2, Tbx21,* and *Ifng*). Impaired UTX function has significant functional impact, since T cell-specific UTX deletion protects mice from the development of colitis. Together, these data elucidate a previously unappreciated hypoxia-associated adaptive response in which impairment of UTX‘s H3K27me3 demethylase activity attenuates CD4^+^ T cell pathogenicity in colitis.

## Results

### Hypoxia-associated decrease in CD4^+^ T cell IFN-γ production is recapitulated by UTX deficiency

In multiple studies, hypoxia has been associated with decreased CD4^+^ T cell IFN-γ production^10,12,13,18^, and our analyses of mouse CD4^+^ T cells corroborate these findings. CD4^+^ T cells stimulated with proinflammatory cytokines IL-2 and IL-12 produced lower levels of IFN-γ in hypoxic (1% O_2_), compared to normoxic (20% O_2_), conditions **(Fig. 1a-c)**. The molecular mechanisms important in mediating these hypoxia-induced changes, however, are not entirely known.

**Figure 1:**
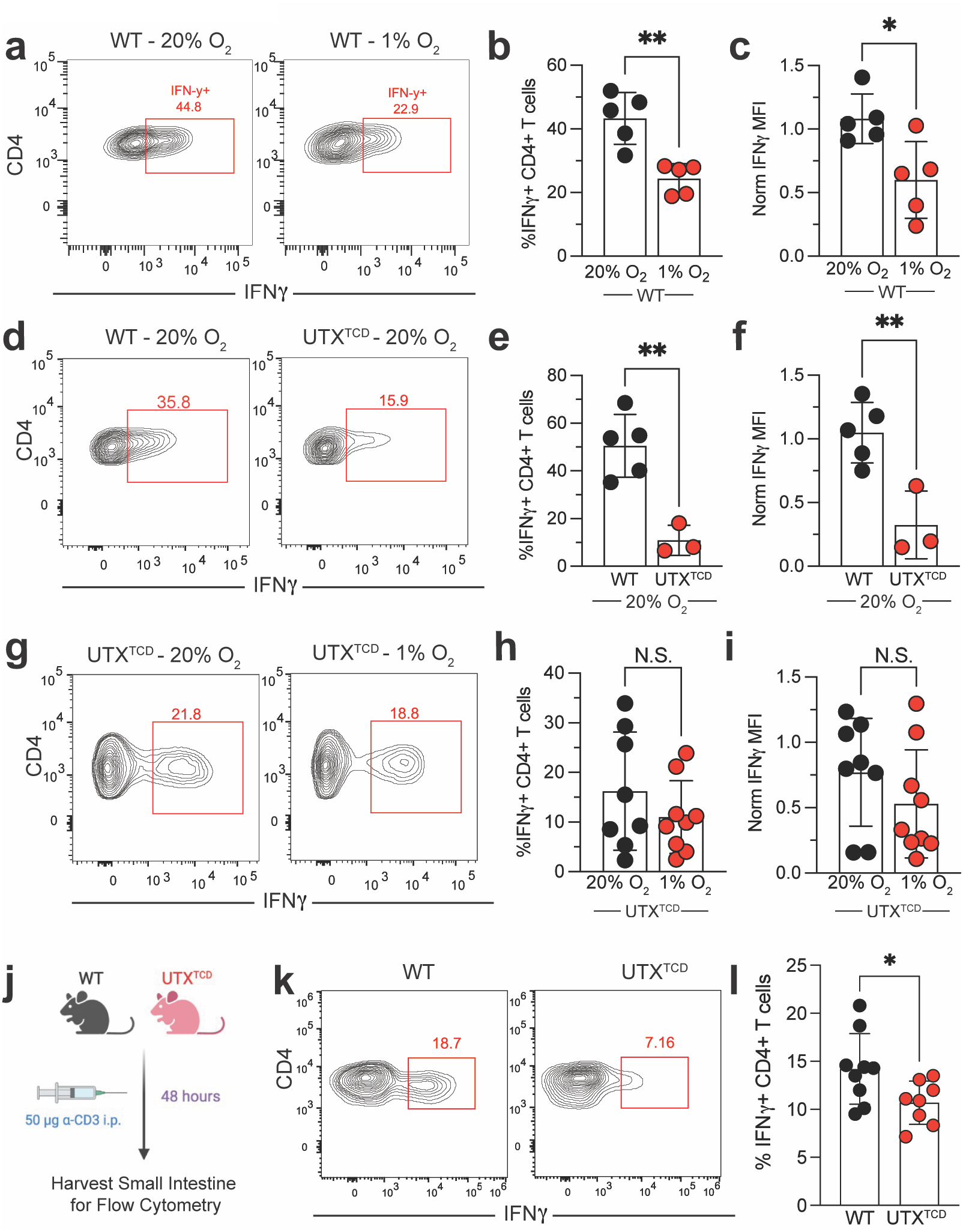
Attenuated CD4^+^ T cell IFN-γ production in hypoxia is recapitulated by UTX deficiency. **a - i)** Flow cytometric analysis of CD4^+^ T cells isolated from spleens of WT and/or UTX^TCD^ mice and stimulated in the presence of IL-12 (10 ng/ml), IL-2 (5ng/ml), anti-CD28 (0.5 mg/ml), and plate-bound anti-CD3 (10 ug/ml) for 4 days. **a)** Representative contour plots and quantification of **b)** %IFN-γ^+^ and **c)** IFN-γ MFI in WT CD4^+^ T cells cultured in normoxia (20% O_2_) vs. hypoxia (1% O_2_) (n = 5 per group). **d)** Representative contour plots and quantification of **e)** %IFN-γ^+^ and **f)** IFN-γ MFI in WT vs. UTX^TCD^ CD4^+^ T cells cultured in normoxia (20% O_2_) (WT: n = 5; UTX^TCD^: n = 3). **g)** Representative contour plots and quantification of **h)** %IFN-γ^+^ and **i)** IFN-γ MFI in WT and UTX^TCD^ CD4^+^ T cells cultured in normoxia (20% O_2_) compared to hypoxia (1% O_2_) (UTX^TCD^: n =8). **j)** Schematic of experimental design: WT and UTX^TCD^ mice were injected with 50 ug of anti-CD3 antibody intraperitoneally (i.p.). After 48 hours, the small intestine was harvested for flow cytometry. **k)** Representative contour plots and **l)** quantification of %IFN-γ^+^ CD4^+^ T cells from the small intestine of WT or UTX^TCD^ mice (n = 9 per group). Data are representative of 2-3 independent experiments. Samples were compared using unpaired two-tailed Student’s t test and data points are presented as individual mice with the mean ± SEM (N.S.; Not Significant; *, p<0.05; **, p<0.01).

Recent work has demonstrated that the histone demethylase UTX is a direct oxygen sensor^8^. As a member of the 2-oxoglutarate (OG)-dependent dioxygenase (2-OGDD) family of enzymes, UTX’s demethylase activity is oxygen-dependent and deactivated in hypoxia^8^. Given previous reports that UTX regulates hypoxia-induced pathways in mouse myoblast cells^8^, we hypothesized that UTX may also play a role in hypoxia-induced changes in immune cells. To interrogate UTX’s function in T cells, we produced mice with a T cell specific deletion of UTX (*Kdm6a*^flox/flox^ *Lck^cre^*^+^; hereafter referred to as UTX T cell-deficient or UTX^TCD^ mice). Remarkably, UTX deficiency mirrored the effects of hypoxia, with significantly decreased IFN-γ production by UTX^TCD^ CD4^+^ T cells, compared to littermate *Kdm6a*^flox/flox^ *Lck^cre^*^-^ controls (hereafter referred to as WT) at 20% O_2_ **(Fig. 1d-f)**. Differences seen in CD4^+^ T cell IFN-γ production during normoxia vs. hypoxia were erased by UTX deficiency **(Fig. 1g-i)**, suggesting that UTX is required for CD4^+^ T cell oxygen sensitivity. Together, these data demonstrate that loss of UTX function phenocopies the decreased IFN-γ production in hypoxic conditions and that UTX is required for hypoxia-sensitivity of CD4^+^ T cells.

To determine whether UTX deficiency also attenuates CD4^+^ T cell production of IFN-γ *in vivo*, we utilized a system in which injection with anti-CD3 antibody triggers CD4^+^ T cell-mediated inflammation in the small intestine **(Fig. 1j)**^19^. Compared to WT mice, a lower frequency of IFN-γ production was detected in CD4^+^ T cells in the small intestine of UTX^TCD^ mice **(Fig. 1k, l)**. Thus, UTX deficiency impairs CD4^+^ T cell IFN-γ production *in vitro* as well as *in vivo*.

### UTX deficiency in CD4^+^ T cells is associated with H3K27me3 accumulation at Th1 pathway genes

In the setting of hypoxia, accumulation of H3K27me3 marks has been demonstrated in mouse hepatoma cells, human renal carcinoma cells, and other cell types^8^. In parallel, our data show H3K27me3 accumulation after incubation in hypoxic (1% O_2_) vs. normoxic (20% O_2_) oxygen conditions in primary mouse CD4^+^ T cells **(Fig. 2a)**. Thus, hypoxia is associated with increased H3K27me3 marks across multiple cell types, including T cells.

**Figure 2:**
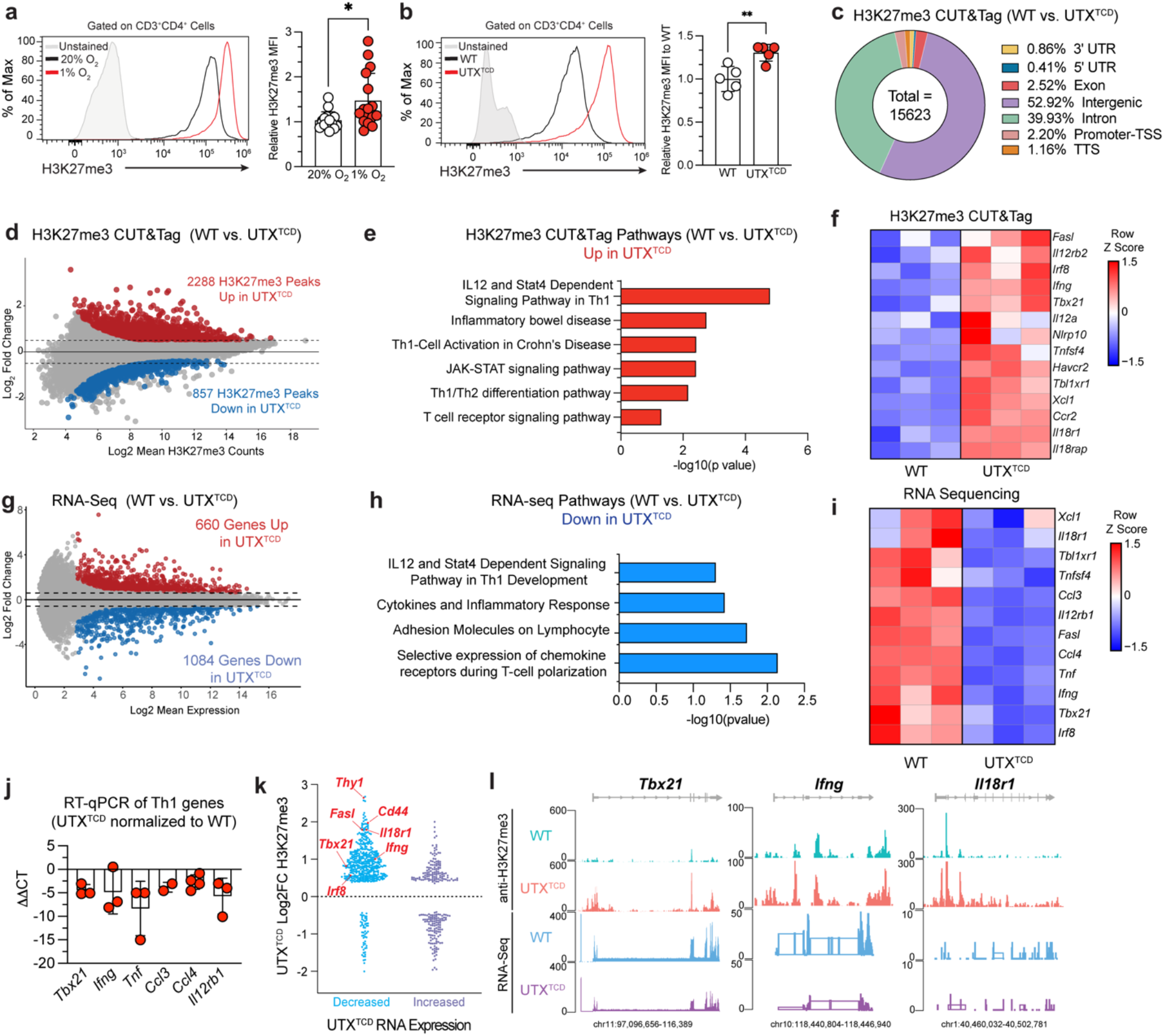
UTX promotes CD4^+^ T cell inflammatory pathways through demethylation of Th1 genes. **a-b)** Flow cytometric analysis of CD4^+^ T cells isolated from spleens of WT and/or UTX^TCD^ mice that were stimulated with anti-CD28 (0.5 mg/ml) and plate-bound anti-CD3 (10 ug/ml) for 48 hours. **a)** Representative histograms (left) and quantification (right) of H3K27me3 MFI in CD4^+^ T cells in normoxia (20% O_2_) vs. hypoxia (1% O_2_) (n = 15 per group). **b)** Representative histograms (left) and quantification (right) of H3K27me3 MFI in CD4^+^ T cells of WT vs. UTX^TCD^ mice (n = 5 per group) in normoxia (20% O_2_). **(c-f)** H3K27me3 CUT&Tag of WT and UTX^TCD^ CD4^+^ T cells stimulated with anti-CD28 (0.5 mg/ml), and plate-bound anti-CD3 (10 ug/ml) for 48 hours. **c)** Pie chart indicating location of differential H3K27me3 marks between WT vs. UTX^TCD^ CD4^+^ T cells. **d)** Volcano plot of differential H3K27me3 marks in WT vs. UTX^TCD^ CD4^+^ T cells plotted as Log_2_ Fold Change (y-axis; Log_2_ Fold Change > 0.5) and Log_2_ Mean H3K27me3 Counts (x-axis; p-value < 0.05). Red dots indicate peaks that are significantly increased (Log_2_ Fold Change > 0.5 and p-value < 0.05) and blue dots indicate peaks that are significantly decreased (Log_2_ Fold Change > - 0.5 and p-value < 0.05). Gray dots are not significant (p > 0.05). **e)** Pathway analysis of loci with significantly increased H3K27me3 using g:profiler with -log_10_(p-value) plotted on x-axis. **f)** Heatmap of target genes in Th1 gene pathways that display differential H3K27me3 marks. **g-i)** Bulk RNA-seq of WT or UTX^TCD^ CD4^+^ T cells after 48 hours of stimulation with anti-CD28 (0.5 mg/ml) and plate-bound anti-CD3 (10 ug/ml) (n = 3 per group). **g)** Volcano plot of differentially expressed genes in WT vs. UTX^TCD^ CD4^+^ T cells plotted with Log_2_ Fold Change (y-axis; Log_2_Fold Change > 0.5) and Log_2_ Mean Expression (x-axis; p-value<0.05). Red dots indicate peaks that are significantly increased (Log_2_ Fold Change > 0.5 and p-value < 0.05) and blue dots indicate peaks that are significantly decreased (Log_2_ Fold Change > −0.5 and p-value < 0.05). Gray dots are not significant (p > 0.05). **h)** Pathway analysis of significantly downregulated RNA-seq genes using DAVID with -log_10_(p-value) plotted on x-axis. **i)** Heatmap of target genes in Th1 gene pathways that are significantly differentially expressed by RNA-seq in WT or UTX^TCD^ CD4^+^ T cells. **j)** qRT-PCR of relative Th1 gene expression in UTX^TCD^ CD4^+^ T cells normalized to WT. **k)** Log_2_ Fold Change (FC) in UTX^TCD^ vs. WT of H3K27me3 deposition (y axis) plotted by either decreased (>-0.5 Log2FC) (blue) or increased (>+0.5 Log2FC) (purple) expression by RNA-seq (x-axis). **l)** Representative gene tracks from UCSC Integrated Genome Browser of H3K27me3 CUT&Tag and RNA sequencing of *Ifng*, *Tbx21*, and *Il18r* in WT and UTX^TCD^ CD4^+^ T cells; Y-axis depicts counts per million (CPM). **a,b)** Data are representative of 2-3 independent experiments. Samples were compared using unpaired two-tailed Student’s t test and data points are presented as individual mice with the mean ± SEM (N.S.; Not Significant; *, p<0.05; **, p<0.01).

As a H3K27me3 demethylase, UTX catalyzes the removal of H3K27me3 histone marks. Loss of UTX’s demethylase activity, then, would be predicted to result in accumulation of H3K27me3 marks **(Supp. Fig. 1a).** Notably, hypoxia has recently been reported to impair UTX’s demethylase activity^8^. Thus, we hypothesized that the observed H3K27me3 accumulation in T cells in low oxygen conditions is due to hypoxia-induced impairment of UTX’s demethylase function. In support of this hypothesis, H3K27me3 levels were higher in CD4^+^ T cells from UTX^TCD^ mice compared to WT **(Fig. 2b)**. Moreover, mouse CD4^+^ T cells treated with a H3K27me3 inhibitor (GSKJ4)^20^ also increased H3K27me3 deposition **(Supp. Fig. 1b, c)**. These findings support a model in which T cells accumulate H3K27me3 during hypoxia through hypoxia-induced impairment of UTX’s demethylase function.

Previously, naïve CD4^+^ T cell differentiation to IFN-γ producing T helper type 1 (Th1) cells was associated with removal of repressive H3K27me3 marks at *Ifng, Tbx21,* and other Th1-associated loci^21^. Erasure of these repressive H3K27me3 marks increases chromatin accessibility at these genetic loci and poises them for active gene transcription to initiate Th1 differentiation. Which epigenetic regulator is responsible for removal of these H3K27me3 marks, however, is not clear.

Since UTX deficiency in T cells results in decreased IFN-γ production by CD4^+^ T cells, we reasoned that removal of H3K27me3 at Th1 loci may be catalyzed by UTX. To identify specific genetic loci regulated by UTX’s H3K27me3 demethylase activity, we performed concomitant H3K27me3 CUT&Tag and RNA sequencing in splenic CD4^+^ T cells isolated from UTX^TCD^ vs. WT mice. H3K27me3 CUT&Tag showed that the majority of sequenced reads mapped to enhancer-associated intergenic (52.92%) and intronic (39.93%) regions, with a small subset in promoter-TSS (transcription start site) regions (2.2%) **(Fig. 2c)**. Additionally, H3K27me3 accumulation was evident in UTX^TCD^ CD4+ T cells compared to WT, with increased H3K27me3 deposition at 2288 peaks vs. decreased deposition at 867 peaks (adjusted p-value <0.05; FDR<0.05; FC>±0.5) **(Fig. 2d)**. Gene pathway ontology analysis of H3K27me3-associated gene loci revealed enrichment of multiple inflammatory pathways (“JAK-STAT signaling”, “T cell receptor signaling”) and Th1 associated pathways (e.g. “IL12 and Stat4 Dependent signaling pathway in Th1”, “Th1/Th2 differentiation pathway”) in UTX^TCD^ CD4^+^ T cells **(Fig. 2e).** Specifically, increased H3K27me3 deposition was observed at key Th1-associated genes (e.g. *Ifng, Tbx21, Il18r1*) in UTX^TCD^ CD4^+^ T cells, compared to WT **(Fig. 2f).** Together, these data demonstrate that UTX catalyzes removal of repressive H3K27me3 marks at multiple Th1-associated gene loci in CD4^+^ T cells.

At the same time, bulk RNA sequencing revealed global changes in transcription, with 1084 downregulated and 660 upregulated genes (adjusted p-value <0.05; FDR<0.05; FC>±0.5) in UTX^TCD^ splenic CD4^+^ T cells, compared to WT **(Fig. 2g)**. Gene ontology pathway analysis of genes downregulated in UTX^TCD^ CD4^+^ T cells showed enrichment of “IL-12 and Stat4 dependent signaling,” “Pathways of Th1 Development”, and “Cytokines and Inflammatory Response” **(Fig. 2h)**. Downregulated genes in UTX^TCD^ CD4^+^ T cells included multiple Th1 genes (*Tbx21, Ifng, Ccl3, Irf8, Tnf,* and *Il18r1)* **(Fig. 2i).** Transcriptional repression of multiple Th1 genes in UTX^TCD^ CD4^+^ T cells was verified by qRT-PCR **(Fig. 2j).** These data implicate UTX in activating Th1 gene transcription in CD4^+^ T cells.

We next sought to identify Th1 genes whose expression are directly activated by UTX’s H3K27me3 demethylase activity. Loss of UTX’s demethylase activity is predicted to decreased gene transcription through accumulation of repressive H3K27me3 marks at target gene loci **(Supp. Fig. 1a)**. Remarkably, multiple genes downregulated in their transcriptional activity in UTX-deficient CD4+T cells also show accumulation of repressive H3K27me3 marks (e.g., *Thy1, Cd44, Ifng, Il18r1, Tbx21, Irf8*, and *Fasl*) **(Fig. 2k)**. For instance, significantly higher H3K27me3 accumulation was seen at loci associated with *Tbx21, Ifng,* and *Il18r1* by H3K27me3 CUT&Tag in UTX^TCD^ vs. WT CD4^+^ T cells **(Fig. 2l**, top two gene tracks), which was accompanied by concomitant decreased transcription at these gene loci **(Fig. 2l**, bottom two gene tracks). Together, these findings suggest that UTX’s H3K27me3 demethylase activity directly catalyzes the removal of repressive H3K27me3 marks to activate transcription at Th1 loci.

### Hypoxia and T cell specific UTX deficiency is associated with increased Tregs

Previous work has demonstrated that hypoxia not only impairs IFN-γ production by CD4^+^ T cells, but also increases generation of suppressive CD4^+^ CD25^+^ FOXP3^+^ T regulatory cells (Tregs)^13,17^. We confirmed these findings using mouse primary naive CD4^+^ T cells after 2 days in culture with anti-CD3 and anti-CD28 in hypoxic (1% O_2_) vs. normoxic (20% O_2_) conditions **(Fig. 3a)**. In parallel, we noted an increased frequency of Tregs in CD4^+^ T cells isolated from UTX^TCD^ mice, compared to WT littermates **(Fig. 3b).** Additionally, after anti-CD3 antibody injection to induce small intestine inflammation, Treg frequencies were increased in the lamina propria (LP), mesenteric lymph nodes (MLN), small intestine (SI), and spleen of UTX^TCD^ mice, compared to WT littermates **(Fig. 3c,d)**. Together, these findings demonstrate that increased Treg frequency occurs in both hypoxia and UTX deficiency, suggesting that UTX impairment may be responsible for hypoxia-induced Treg differentiation.

**Figure 3:**
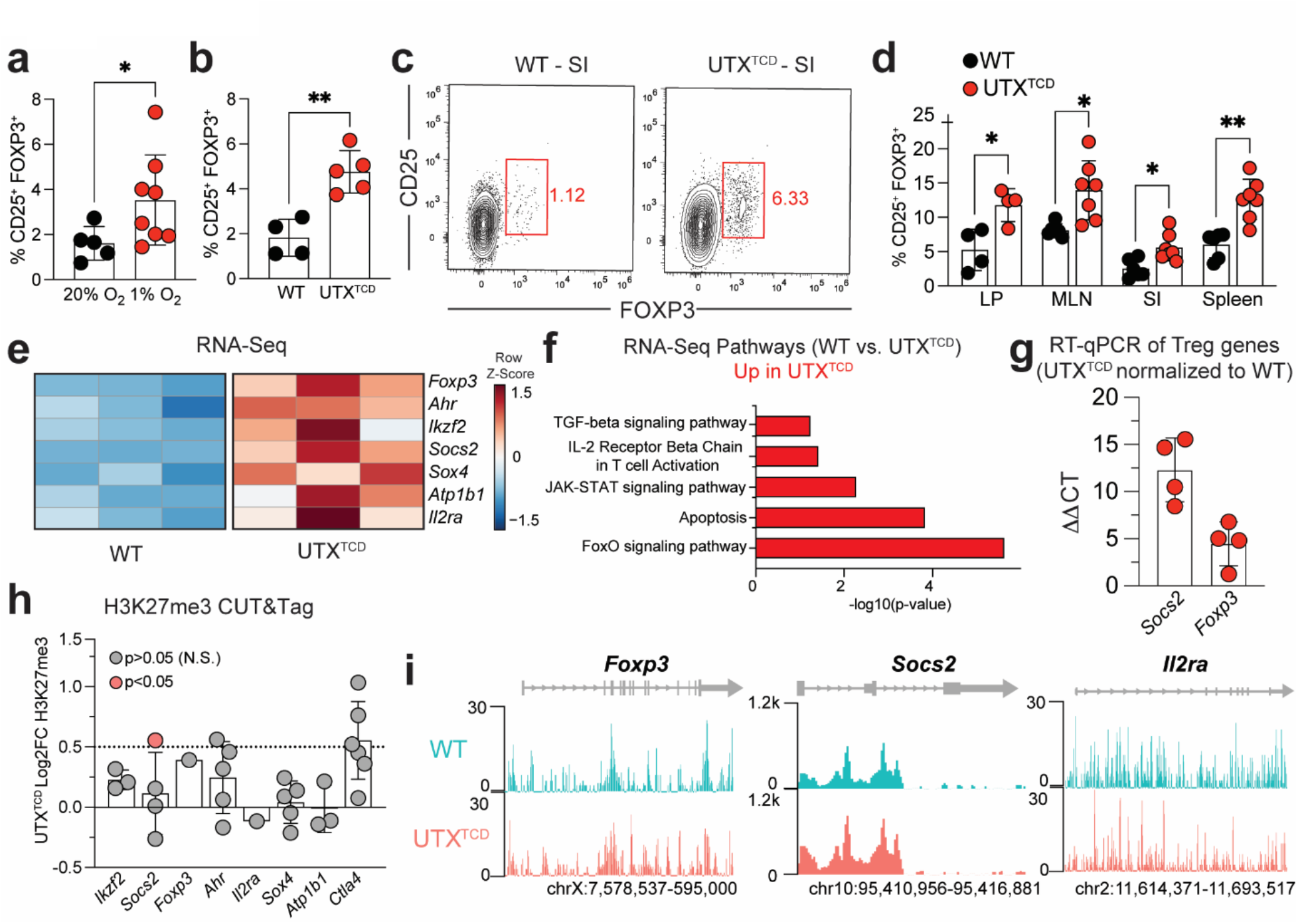
Hypoxia and UTX deficiency are both associated with increased Tregs. **a-b)** Flow cytometric analysis of splenic CD4^+^ T cells from WT and/or UTX^TCD^ mice after stimulation with anti-CD28 (0.5 mg/ml) and plate-bound anti-CD3 (10 ug/ml) for 48 hours. **a)** Quantification of %CD25^+^FOXP3^+^ Tregs among CD4^+^ T cells in normoxia (20% O_2_; n=5) vs. hypoxia (1% O_2_; n = 9). **b)** Quantification of CD25^+^FOXP3^+^ among CD4^+^ T cells in WT (n=4) vs. UTX^TCD^ (n=5) mice in normoxia. **c, d)** WT and UTX^TCD^ mice were injected with 50 ug of anti-CD3 antibody intraperitoneally (i.p.). After 48 hours, the lamina propria (LP), mesenteric lymph nodes (MLN), small intestine (SI), and spleen were harvested for flow cytometry. **c)** Representative flow cytometric analysis of CD4^+^ T cells from small intestine (SI). **d)** Quantification of %CD25^+^FOXP3^+^ Tregs among CD4^+^ T cells in organs shown. **e)** Heatmap of differentially expressed genes (p-value<0.05) by bulk RNA-seq of WT vs. UTX^TCD^ splenic CD4^+^ T cells after 48 hours of stimulation with anti-CD28 (0.5 mg/ml) and plate-bound anti-CD3 (10 ug/ml) (n=3 per group), as described in Figure 2. **f)** Pathway analysis of significantly downregulated RNA-seq genes using DAVID with -log_10_(p-value) plotted on x-axis. **g)** qRT-PCR of Treg genes in UTX^TCD^ CD4^+^ T cells normalized to WT. **h and i)** H3K27me3 CUT&Tag analysis of WT and UTX^TCD^ CD4^+^ T cells stimulated with anti-CD28 (0.5 mg/ml), and plate-bound anti-CD3 (10 ug/ml) for 48 hours, as shown in Figure 2. **h)** Log_2_Fold Change (FC) of Treg pathway genes from H3K27me3 CUT&Tag, with each point indicating an individual peak associated with that gene. Red circle indicates significant p-values<0.05. Grey circle indicates peaks that are not significant (N.S.; p>0.05). **i)** Representative gene tracks from UCSC Integrated Genome Browser of H3K27me3 CUT&Tag of *Foxp3*, *Socs2*, and *Il2ra*; Y-axis depicts counts per million (CPM). **a-d)** Data are representative of 2-3 independent experiments. Samples were compared using unpaired two-tailed Student’s t test and data points are presented as individual mice with the mean ± SEM (N.S.; Not Significant; *, p<0.05; **, p<0.01).

On bulk RNA sequencing, expression of Treg-associated genes (*Foxp3, Ahr, Ikzf2, Soc2, Sox4, Atp1b1, Il2ra*) were significantly higher in UTX^TCD^ CD4^+^ T cells, compared to WT **(Fig. 3e).** Gene ontology analysis showed upregulation of multiple Treg-associated pathways (e.g. “IL-2 Receptor Βeta Chain in T cell activation”, “TGF-β signaling”) **(Fig. 3f)**. Transcriptional upregulation of *Socs2* and *Foxp3* in UTX^TCD^ CD4^+^ T cells was verified by qRT-PCR (real-time quantitative reverse transcription polymerase chain reaction) **(Fig. 3g)**. In contrast to Th1 genes, however, Treg genes are unlikely to be the direct targets of UTX’s demethylase activity. Because UTX removes repressive H3K27me3 marks, UTX deficiency would be predicted to repress, rather than enhance, gene transcription **(Supp. Fig. 1a)**. Moreover, no significant differences in H3K27me3 deposition were seen at *Ikzf2, Foxp3, Ahr, Il2ra, Sox4*, *Atp1b1*, or *Ctla4* in WT vs. UTX^TCD^ CD4^+^ T cells **(Fig. 3h, 3i)**, suggesting that Treg loci are not direct targets of UTX’s demethylase activity. Instead, UTX deficiency may result in increased Treg gene transcription through an indirect mechanism. Because *Ifng* and *Tbx21* expression antagonizes Treg differentiation^22,23^, for instance, decreased *Ifng* and *Tbx21* expression in UTX-deficient T cells may indirectly promote expression of Treg genes.

### T cell specific UTX deficiency protects against colitis

Multiple lines of evidence demonstrate a key role for immune dysregulation in the development of IBD. Strong evidence exists that increased T cell IFN-γ production is linked to IBD development, and CD4^+^ T cell IFN-γ deficiency is protective in mouse models of colitis^14–16^. Additionally, decreased Treg numbers and immunosuppressive function are linked to IBD, and Treg transfer can ameliorate colitis development in mouse models^24–26^. Based on our data that UTX deficiency lowers CD4^+^ T cell IFN-γ production and increases Treg numbers, we hypothesized that impairment of T cell UTX may restore immune homeostasis in IBD and protect against colitis.

To test this, we utilized a CD4^+^ T cell adoptive transfer model of inflammatory colitis in which naïve CD4^+^ T cells (CD3^+^ TCRβ^+^ CD4^+^ CD45RB^hi^) from either WT or UTX^TCD^ mice were transferred into immunodeficient recipients (Rag2^-/-^ mice) **(Fig. 4a)**^27^. Notably, recipients of UTX^TCD^ CD4^+^ T cells were protected from pathologies associated with colitis, including weight loss and lethality (defined by >20% weight loss) **(Fig. 4b, c)**. Hematoxylin and Eosin staining of colons revealed diffuse inflammation from the cecum to rectum with immune infiltration into the lamina propria in the recipients of WT cells **(Fig. 4d, left, and Fig. 4e)**. In contrast, recipients of UTX^TCD^ T cells had significantly decreased lymphocytic infiltration into the lamina propria, mucosal thickening, transmural spread, and abnormal crypt formation as measured by blinded histopathological scoring **(Fig. 4d, right, and Fig. 4e)**. Moreover, recipients of UTX^TCD^ CD4^+^ T cells had decreased colon inflammation, as measured by colon weight to length ratios **(Fig. 4f)** and decreased IFN-γ production by CD4^+^ T cells compared to WT in the lamina propria **(Fig. 4g)**. Additionally, transfer of UTX^TCD^ CD4^+^ T cells results in an increased proportion of Tregs in the lamina propria **(Fig. 4h)**. Together, these data demonstrate that CD4^+^ T cell-intrinsic loss of UTX function protects against inflammatory colitis through suppressed IFN-γ production and increased Treg numbers.

**Figure 4:**
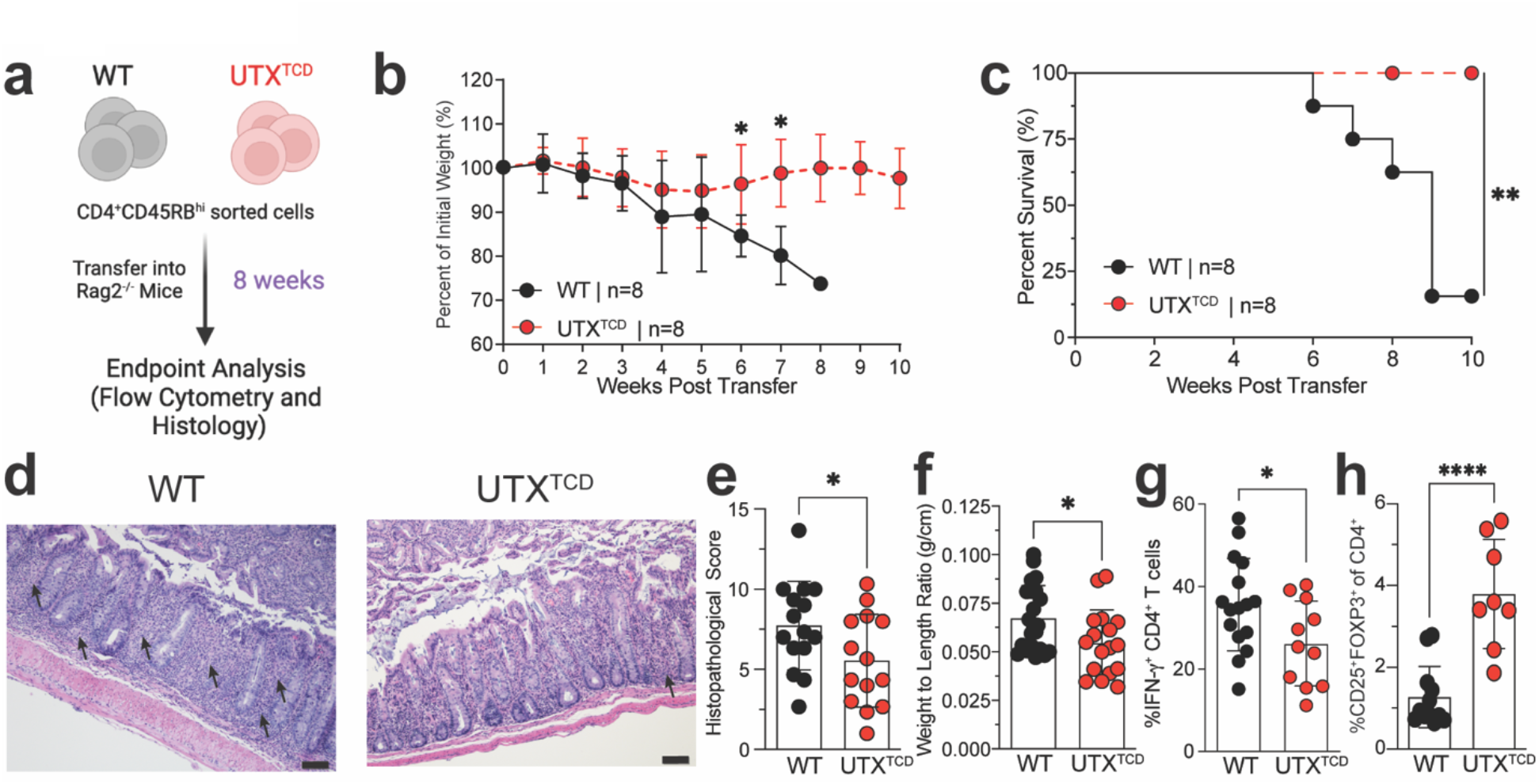
T cell UTX deficiency protects against colitis. **A)** Schematic of experimental design: Sorted WT I. UTX^TCD^ CD4^+^ CD45RB^hi^ T cells were transferred into Rag2^-/-^ recipients. Recipients were monitored for 8 weeks and then sacrificed for flow cytometry and histological analysis of the lamina propria. **b)** Percent weight loss from original weight in recipients of WT vs. UTX^TCD^ transferred cells (n = 7-8 per group). **c)** Kaplan-Meier survival curve of recipients of WT vs. UTX^TCD^ transferred cells (n = 7-8 per group). **d)** Representative Hematoxylin and Eosin (H&E) staining of colon tissues at study endpoint. Arrows denote extensive infiltration in the lamina propria in recipients of WT transferred cells (left), compared with mild infiltration in recipients of UTX^TCD^ transferred cells (right). Scale bar=50um. **e)** Average histopathological scores of colon tissues. **f)** Weight-to-length ratios of colons harvested at study endpoint, measured as weight (g) divided by length (cm) from the cecum to rectum (n = 10 per group). **g,h)** Flow cytometric quantification of %IFN-γ^+^ **(g)** and CD25^+^FOXP3^+^ Treg cells **(h)** among CD4^+^ T cells from the lamina propria of recipients I WT vs. UTX^TCD^ transferred cells (n = 16 per group). Samples were compared using unpaired two-tailed Student’s t and data points are presented as individual mice with the mean ± SEM (*, p<0.05; ****, p<0.0001).

## Discussion

Identifying adaptive mechanisms that protect against hypoxia’s deleterious effects is an important step toward harnessing these pathways as therapeutic strategies for IBD^1,6^. However, the mediators important in these protective responses in IBD remain incompletely defined. Here we identify a mechanism mediated by the oxygen-sensitive epigenetic regulator UTX that is protective in an inflammatory colitis model **(Supp. Fig. 2)**. Our results corroborate previous findings that hypoxia is associated with decreased CD4^+^ T cell IFN-γ production and increased Tregs. Notably, T cell-specific UTX deficiency recapitulates both of these findings. Furthermore, both hypoxia and UTX-deficiency results in H3K27me3 accumulation, confirming previous reports that hypoxia impairs the H3K27me3 demethylase activity of UTX^8,9^. Concomitant RNA-seq and H3K27me3 CUT&Tag identified multiple Th1 genes (*Tbx21, Ifng,* and *Il18r1*) as direct targets of UTX’s histone demethylase activity. Together, these findings demonstrate a T cell-intrinsic role for UTX’s histone demethylase activity, which is oxygen-sensitive, in modulating severity of colonic inflammation.

While our data implicate UTX’s demethylase activity in the direct regulation of Th1 genes, how UTX controls Treg gene expression is more complex. Our results show increased Treg gene expression in UTX-deficient CD4^+^ T cells, which do not correlate with H3K27me3 changes at those gene loci. These findings suggest that UTX does not directly target Treg loci for H3K27me3 demethylation, but instead regulates Treg numbers in an indirect manner. One possibility is that increased Treg numbers are due to decreased expression of *Ifng* and *Tbx21*, which antagonize Treg differentiation^22,23^. Furthermore, UTX can also cooperate with additional epigenetic regulators such as BRG1, MLL4/5, SWI/SNF to deposit additional histone marks and modulate general chromatin accessibility in a demethylase-independent manner^28–30^. Whether these additional UTX-mediated demethylase-independent functions are altered in hypoxia remains to be determined.

Because UTX is ubiquitously expressed, hypoxia may also alter UTX-dependent gene expression across multiple cell types, including diverse immune cell types found at the mucosal barrier in IBD. Which immune cell types are altered by hypoxia in an UTX-dependent manner, however, remain unclear. Interestingly, hypoxia has been reported to promote polarization of macrophages to anti-inflammatory M2 state in inflammation^31^. At the same time, loss of UTX in macrophages also induces M2 macrophage polarization^32^. While these findings suggest the possibility that UTX may be responsible for the effects of hypoxia in macrophages, it remains to be tested whether hypoxia may polarize macrophages by impeding UTX’s demethylase activity. Moreover, pharmacological inhibition of UTX H3K27me3 demethylase activity has recently been shown to decrease IFN-γ production in human NK (natural killer) cells^7^. Additional studies are needed to identify the contributions of UTX in NK cells and other immune cell subtypes during mucosal inflammation.

High expression of IFN-γ and IL-17 have been reported in the intestinal mucosa of IBD patients^33,34^. While the pathogenic role of IFN-γ in mouse models of IBD is clear^15,35^, the role of IL17 is more complex. While some studies suggest that IL-17-producing T helper 17 (Th17) cells promote IBD pathogenesis^36,37^, other studies have reported that IL-17 is protective^38,39^. Interestingly, hypoxia has been reported to promote *in vitro* Th17 cell differentiation from naïve CD4^+^ T cells^40^. However, additional studies will be required to understand the role of IL-17 and Th17 cells in adaptive responses to hypoxic inflammation.

To date, efforts to harness adaptive hypoxia pathways have targeted pathways important in protecting mucosal barrier function. For example, GB004 has shown efficacy in early clinical trials, and GB004-mediated HIF stabilization is proposed to be disease-protective through its effects on the gut epithelium^1,6^. Importantly, the pathogenesis of IBD involves both the breakdown of mucosal epithelial barrier function as well as the dysregulation of the mucosal immune response. In addition to architectural disturbances of mucosal surfaces, histopathological examination of biopsies from patients with active disease demonstrate areas of large immune cell infiltrate^16,41^. Here we identify a hypoxia-sensitive pathway mediated by UTX that plays an immune-protective role in colitis. These findings suggest that pharmacological inhibitors of UTX’s H3K27me3 demethylase activity (e.g., GSK-J4)^20^ allow for therapeutic manipulation of UTX-dependent pathways in inflammatory colitis. By targeting a distinct component of pathogenesis, GSKJ4 may have additive benefits in combination with GB004 in the treatment of IBD.

## Acknowledgements

We thank members of the Su lab for helpful discussion. We thank Melissa Lechner for collaborating with BioRender platform to produce graphical figure. M.A.S. is supported by the NIH (NS107851, AI143894, DK119445) Department of Defense (USAMRAA PR200530), and National Organization of Rare Diseases. M.I.C. is supported by Ruth L. Kirschstein National Research Service Awards (GM007185 and AI007323), and Whitcome Fellowship from the Molecular Biology Institute at UCLA.

---

**Supplemental Figure 1:**
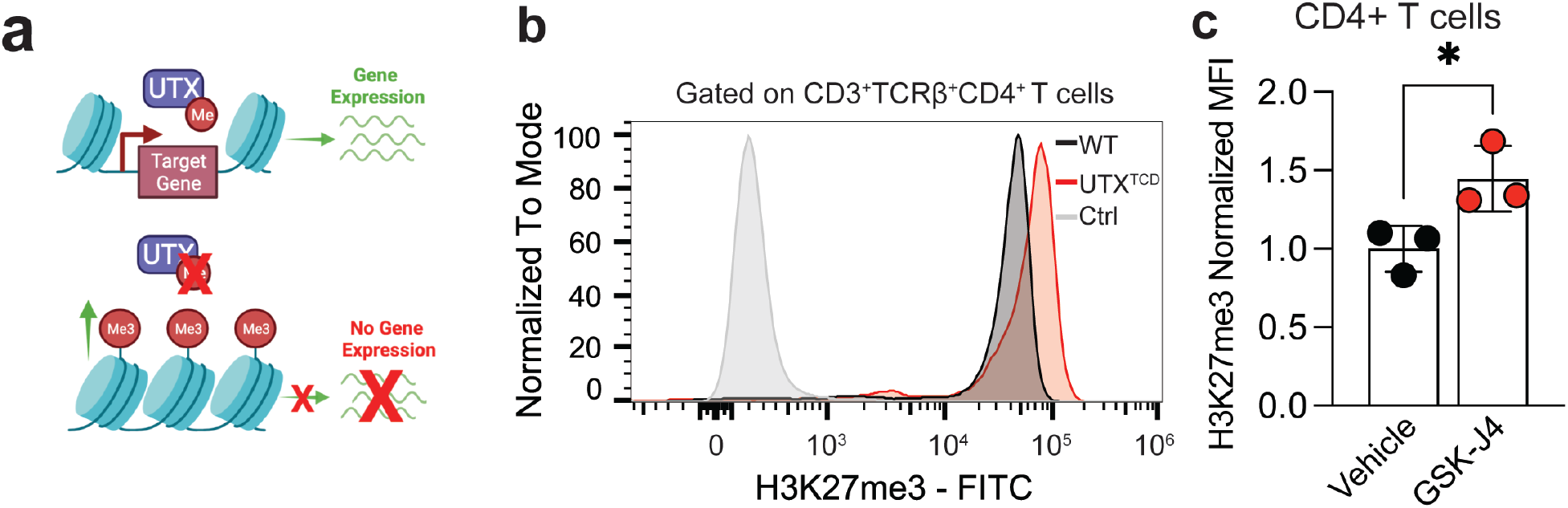
Treatment of GSK-J4 results in accumulation of H3K27me3 in CD4^+^ T cells. **a)** Schematic of UTX’s role as a H3K27me3 demethylase. Top, UTX demethylase activity leads to removal of repressive H3K27me3 marks and activation of target gene expression. Bottom, when UTX demethylase activity is impaired, H3K27me3 accumulates and target gene expression is downregulated. **b, c)** Flow cytometric analysis of splenic CD4^+^ T cells from WT and UTX^TCD^ mice that were stimulated with anti-CD28 (0.5 mg/ml) and plate-bound anti-CD3 (10 ug/ml) and treated with either vehicle control (DMSO) or GSK0J4 (30 uM) for 48 hours. **b)** Representative histograms **c)** and quantification of H3K27me3 MFI in CD4^+^ T cells (n = 3 per group). Samples were compared using unpaired two-tailed Student’s t and data points are presented as individual mice with the mean ± SEM (*, p<0.05).

**Supplemental Figure 2:**
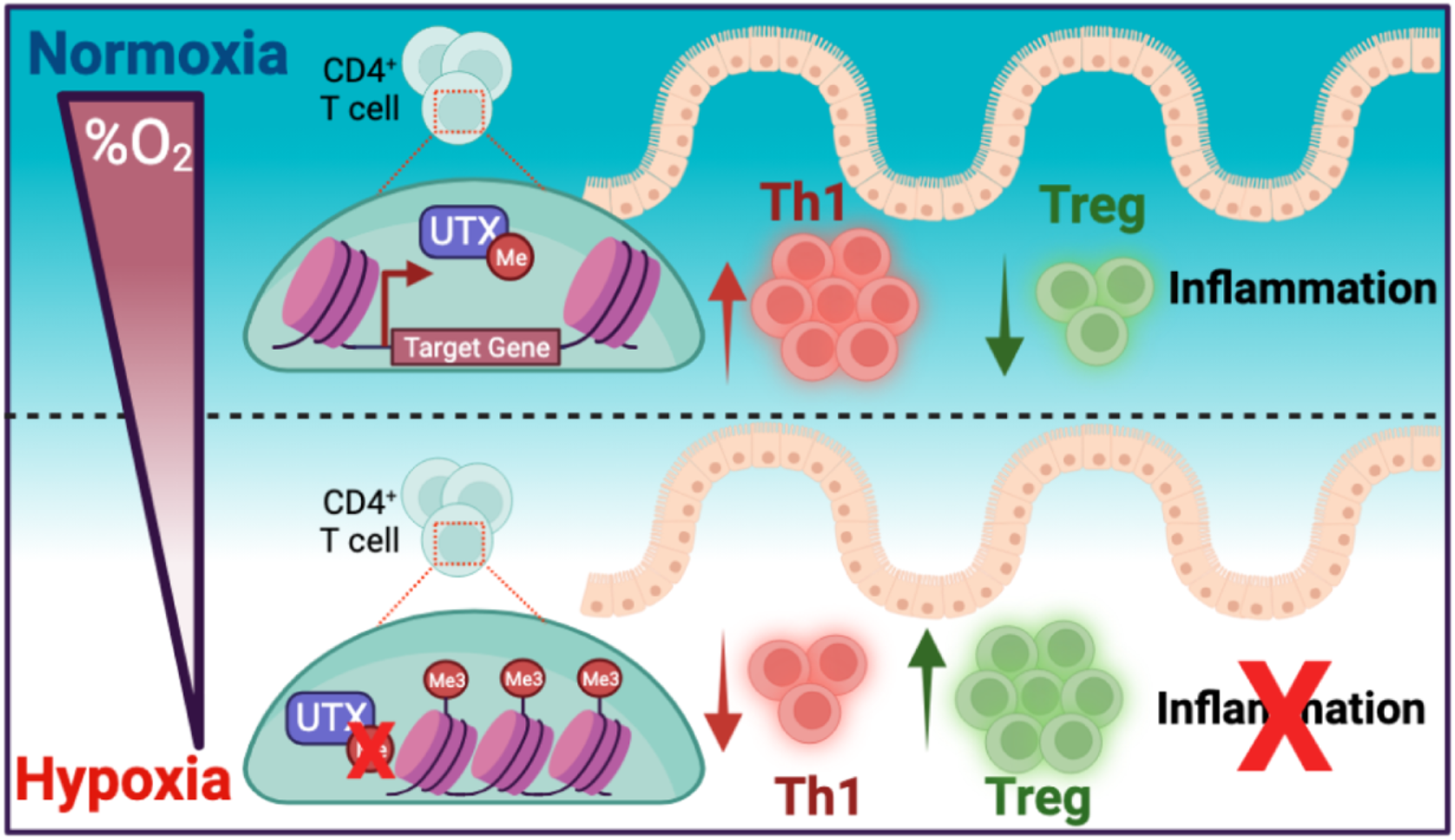
UTX is an oxygen-sensitive master regulator of Th1 and Treg gene programs in autoimmunity. In normoxia, UTX demethylase activity is active, resulting in demethylation of H3K27me3 levels at target genes which promote increased Th1 and decreased Treg populations, resulting in enhanced proinflammatory conditions which promote autoimmunity. However, in hypoxic conditions in which UTX demethylase is inactive, decreased proinflammatory Th1 and increased immunosuppressive Treg populations result in protection from autoimmunity.

## Materials and Methods

### Mice

All mouse experimental procedures were approved by the University of California, Los Angeles, and University of North Carolina at Chapel Hill Institutional Animal Care and Use Committee (IACUC) and mice were housed and bred in sterile, specific pathogen-free mouse facilities. Generation of UTX^TCD^ mice: *Kdm6a^flox/flox^ Lck^Cre^* mouse strain was generated as previously described^42^. Age-matched littermate controls (*Kdm6a^flox/flox^* or WT) were used. Rag2^-/-^ mice were acquired through Jackson Laboratories (Strain# 008449; B6.Cg-Rag2tm1.1Cgn/J). For all mouse experiments, female 6-8 week old age-matched littermates were used in accordance with approved institutional protocols.

### T cell isolation and stimulation

Splenic CD4^+^CD25^-^ naïve T cells of WT or UTX^TCD^ mice were sorted per manufacturer’s instructions (StemCell: Cat. No. 19765) and cultured with anti-CD28 (0.5 mg/ml), and plate-bound anti-CD3 (5ug/ml) for 2 days. *In vivo* T cell activation model: WT or UTX^TCD^ mice were injected with 50 ug of purified anti-CD3–(clone - 2C11) for 2 days. Then the small intestine is isolated and harvested for clow cytometry as previously described^19^.

### Experimental colitis and tissue harvesting

Mouse model of colitis: CD4^+^ cells were enriched from *Kdm6a^flox/flox^* (WT control) or *Kdm6a^flox/flox^ Lck^Cre^* (UTX^TCD^) 8- to 12-week-old female donor spleens using the CD4^+^ T cell Isolation Kit II (Miltenyi Biotec) according to manufacturer instructions. Cells were labeled and sorted for CD4^+^CD25^−^CD45RB^hi^ cells by fluorescence-activated cell sorting. Cells (5 × 10^5^) were injected i.p. into 4- to 7-week-old Rag2^−/−^ recipient female mice. Weights were measured weekly from initial cell transfer. Lack of survival was defined as death or weight loss > 20%. Recipient mice were euthanized once weight loss was >20% of initial body weight or at 10 weeks after adoptive transfer and spleens, mesenteric lymph nodes (MLNs), and colons were harvested. Spleens and MLNs were macerated and filtered into a single cell suspension using a 10-mL syringe using 40um cell strainers (BD), followed by incubation with ACK Lysis Buffer (Gibco). Colons were digested in collagenase/dispase and DNase (Roche), followed by Ficoll gradient centrifugation. Lymphocytes were collected and used for flow cytometry analysis.

### Colon histopathology

Colons were harvested at 10 weeks after adoptive transfer or earlier if weight loss of 20% was noted. Specimens were fixed in 10% formalin, sectioned (5 μm), and stained with H&E. Scoring for inflammation was performed as described^41,43^ in a blinded fashion by a veterinary pathologist.

### Flow cytometry and cell sorting

Cells were analyzed for cell surface markers using fluorophore-conjugated antibodies (BioLegend, eBioscience). Cell surface staining was performed in FACS Buffer (2% FBS and 2 mM EDTA in PBS) and intracellular staining was performed by fixing and permeabilizing using the eBioscience Foxp3/Transcription Factor kit for intranuclear proteins or BD Cytofix/Cytoperm kits for cytokines. Flow cytometry was performed using the Attune NxT Acoustic Focusing cytometer (Thermo) and data were analyzed with FlowJo v10.7.2 software (TreeStar). Cell surface and intracellular staining was performed using the following fluorophore-conjugated antibodies: TCRβ (H57-597), CD3 (17A2), CD4 (GK1.5), IFN-γ (XMG1.2), UTX– (N2C1 - GeneTex), Goat anti-rabbit H&L–(Abcam - ab6717), FOXP3 (Bi–legend - FJK-16s), CD25 (Bio–legend - 3C7). Isolated splenic T cells were sorted using Aria-H Cytometer (BD) to >95% purity.

### Hypoxia and Th1 skewing assay

Splenic CD4^+^CD25^-^ naïve T cells of WT or UTX^TCD^ mice were sorted per manufacturer’s instructions (StemCell) and cultured with cytokines (10 ng/mL IL-2 and 5 ng/ml of IL-2), anti-CD28 (0.5 mg/ml), and plate-bound anti-CD3 (5ug/ml) for 4 days. Cells were moved out of anti-CD3 and anti-CD28 into a new well with fresh media and cytokines on day 2. IFN-γ^+^ CD4^+^ T cells were accessed using intracellular cytokine staining on day 4. Cells were either incubated in normoxia (20% O_2_) or hypoxia (1% O_2_) conditions. For normoxia condition, cells were placed in a 37 degree C 5% CO_2_ incubator in normal oxygen conditions. For hypoxia condition, cells were placed in a modular hypoxia incubator chamber (Billups Rothenburg; MIC-101). Chamber was purged; injected with a mixture of 1% O_2_, 5% CO_2_ balanced with nitrogen for 5 minutes at 20 liters/min; and placed in 37 degree C incubator, per manufacturer instructions.

### Quantitative PCR

For quantitative PCR experiments, up to 5 x 10^5^ cells were isolated using naïve T cell EasySep magnetic Isolation kit (StemCell; Cat. No. 19765). RNA was isolated from cells using Quick-RNA Micro-prep kit (Zymo; Cat. No. R1051). Then, RNA was quantified using a NanoDrop and used up to 1 ug to synthesize cDNA using High-Capacity cDNA Reverse Transcription Kit (ThermoFisher; Cat. No. 4368813). Then 1 ul of undiluted cDNA was with a SybrGreen Real-Time PCR assay in a 386 well plate. Each sample was plated at a minimum of 3 technical replicates per probe. Normalized expression was calculated as follows: 1) β-actin was used as a housekeeping gene and used to normalize the amount of each sample for each probe, then 2) each sample was normalized to WT for each gene.

### RNA-seq library construction

RNA was isolated from 50,000 sort-purified NK cells per sample using RNeasy Mini kit (Qiagen). RNA quality was verified using High Sensitivity RNA Screen Tape and excluded samples with a RINe <6.0. RNA-seq libraries were sequenced using Illumina HighSeq 4000 platform (single end, 50bp).

### CUT&Tag Library Preparation

For anti-H3K27me3 CUT&Tag library preparation, nuclei were isolated with cold nuclear extraction buffer (20 mM HEPES, pH 7.9, 10 mM KCl, 0.1% Triton X-100, 20% glycerol, 0.5 mM spermidine in 1X protease inhibitor buffer) and incubated with activated concanavalin A (ConA) coated magnetic beads (Polysciences - 86057-3) in PCR strip tubes at room temperature for 10 minutes. A 1:100 dilution of primary antibody (anti-UTX Cell Signaling Rabbit mAb #33510 or IgG Isotype Control: Cell Signaling Technology #3900S) in antibody buffer (20 mM HEPES pH 7.5; 150 mM NaCl; 0.5 mM Spermidine; 1X Protease inhibitor cocktail (Roche); 0.05% Digitonin, 2 mM EDTA, 0.1% BSA) was added and nuclei were incubated with primary antibodies overnight at 4C. The next day, the strip tubes were incubated on a magnetic tube holder and supernatants were discarded. Secondary antibody (Guinea Pig anti-Rabbit IgG Fisher Scientific - NBP172763) was added diluted 1:100 in Dig-Wash (20 mM HEPES pH 7.5; 150 mM NaCl; 0.5 mM Spermidine; 1X Protease inhibitor cocktail; 0.05% Digitonin) and nuclei were incubated for 1 hour at room temperature. Nuclei were washed four times in Dig-Wash and then incubated with a 1:20 dilution of pAG-Tn5 adapter complex (EpiCypher) in Dig-300 buffer (1x Protease inhibitor cocktail, 20 mM HEPES pH 7.5, 300 mM NaCl, 0.5 mM spermidine) for 1 hour at room temperature. To stop tagmentation, 25 uL Dig-300 buffer with 10 uL 1 M MgCl, 7.5 uL 0.5 M EDTA, 2.5 uL 10% SDS, and 5 uL 10 mg/mL proteinase K was added to each reaction and incubated at 55 degrees for 1 hour. DNA was extracted by phenol:chloroform:isoamyl alcohol separation. DNA was barcoded and amplified using the following conditions: a PCR mix of 25 uL NEBNext 2X mix, 2 uL each of barcoded forward and reverse 10 uM primers, and 21 uL of extracted DNA was amplified at: 58C for 5 min, 72C for 5 min, 98C for 45 sec, 16x 98C for 15 sec followed by 63C for 10 sec, 72C for 1 min. Amplified DNA libraries were purified by adding 1.3x volume of KAPA pure SPRI beads (Roche) to each sample and incubated for 10 minutes at room temperature. Samples were placed on a magnet and unbound liquid was removed. Beads were rinsed twice with 80% ethanol, and DNA was eluted with 25 uL TE buffer. All individually i7-barcoded libraries were mixed at equal molar proportions for sequencing on an Illumina NovaSeq 6000 sequencer.

### Sequencing Data Analysis

H3K27me3 CUT&Tag fastq files were trimmed to remove low-quality reads and adapters using Cutadapt (version 2.3). The reads were aligned to the reference mouse genome (mm10) with bowtie2 (version 2.2.9). Peak calling was performed with MACS2 (version 2.1.1). Peaks/regions identified as H3K27me3-bound (UTX CUT&Tag) were annotated using the annotatepeaks.pl function from the HOMER analysis package. To determine the distance to the nearest TSS (transcription start site), we used the default settings in annotatepeaks.pl, which utilizes RefSeq transcription start sites to determine the closest TSS. For genomic annotation, we used the “Basic Annotation” output provided by the assignGenomeAnnotation program in annotatePeaks.pl. The TSS was defined from −1kB to +100bp, and TTS (transcription termination site) was defined from −100 bp to +1kB. “Basic Annotation” is based on alignments of RefSeq transcripts to the UCSC hosted mouse genome file (mm10). HTseq (version 0.9.1) was used to count the number of reads that overlap each peak per sample. The peak counts for ATAC-seq were analyzed with DESeq2 (version 1.24.0) to identify differentially accessible genomic regions. Peaks with adjusted p-value < 0.05 were considered significantly differentially accessible. The peak counts for H3K27me3 CUT&Tag were visualized with Integrated Genome Browser (version 9.1.8) using mouse genome 2011. RNA sequencing analysis was carried out by first checking the quality of the reads using FastQC. Then, they were mapped with HISAT2 (version 2.2.1) to the mouse genome (mm10). The counts for each gene were obtained by HTSeq. Differential expression analyses were carried out using DESeq2 (version 1.24.0) with default parameters. Genes with adjusted p value <0.05 were considered significantly differentially expressed. Sequencing depth normalized counts were used to plot the expression values for individual genes.

Pathway analysis of clustered RNA-seq data was performed using DAVID, g:profiler, and Enrichr. Top relevant pathways were selected from KEGG Biological Pathways and Gene Ontology Pathways (Biological Processes and Molecular Function).

### Statistical Analyses

For graphs, data are shown as mean ± SEM, and unless otherwise indicated, statistical differences were evaluated using a Student’s t test. For graphs containing multiple groups, either one-way (one treatment or condition) or two-way (multiple treatments or conditions) ANOVA with Tukey’s correction for multiple comparisons was used as stated. For Kaplan-Meier survival curve, samples were compared using the Log-rank (Mantel-Cox) test with correction for testing multiple hypotheses. A p-value < 0.05 was considered significant. Graphs were produced and statistical analyses were performed using GraphPad Prism and ggplot2 library in R. Spearman Correlation on best fit regression line was performed using ggpubr library in R.

## Data Availability

Sequencing datasets are accessible from the Gene Expression Omnibus (GEO) under accession number: GSE236301 (reviewer access token: otgxeccwzlchjkl). Further information and requests for resources and reagents should be directed and will be fulfilled by the corresponding authors.

## References

1 Colgan, S. P. & Taylor, C. T. Hypoxia: an alarm signal during intestinal inflammation. Nat Rev Gastroenterol Hepatol 7, 281–287, doi:10.1038/nrgastro.2010.39 (2010).

2 Van Welden, S., Selfridge, A. C. & Hindryckx, P. Intestinal hypoxia and hypoxia-induced signalling as therapeutic targets for IBD. Nat Rev Gastroenterol Hepatol 14, 596–611, doi:10.1038/nrgastro.2017.101 (2017).

3 Cummins, E. P., Seeballuck, F., Keely, S. J., Mangan, N. E., Callanan, J. J., Fallon, P. G. & Taylor, C. T. The hydroxylase inhibitor dimethyloxalylglycine is protective in a murine model of colitis. Gastroenterology 134, 156–165, doi:10.1053/j.gastro.2007.10.012 (2008).

4 Gupta, R., Chaudhary, A. R., Shah, B. N., Jadhav, A. V., Zambad, S. P., Gupta, R. C., Deshpande, S., Chauthaiwale, V. & Dutt, C. Therapeutic treatment with a novel hypoxia-inducible factor hydroxylase inhibitor (TRC160334) ameliorates murine colitis. Clin Exp Gastroenterol 7, 13–23, doi:10.2147/CEG.S51923 (2014).

5 Marks, E., Goggins, B. J., Cardona, J., Cole, S., Minahan, K., Mateer, S., Walker, M. M., Shalwitz, R. & Keely, S. Oral delivery of prolyl hydroxylase inhibitor: AKB-4924 promotes localized mucosal healing in a mouse model of colitis. Inflamm Bowel Dis 21, 267–275, doi:10.1097/MIB.0000000000000277 (2015).

6 Karhausen, J., Furuta, G. T., Tomaszewski, J. E., Johnson, R. S., Colgan, S. P. & Haase, V. H. Epithelial hypoxia-inducible factor-1 is protective in murine experimental colitis. J Clin Invest 114, 1098–1106, doi:10.1172/JCI21086 (2004).

7 Danese, S., Levesque, B. G., Feagan, B. G., Jucov, A., Bhandari, B. R., Pai, R. K., Taylor Meadows, K., Kirby, B. J., Bruey, J. M., Olson, A., Osterhout, R., Van Biene, C., Ford, J., Aranda, R., Raghupathi, K. & Sandborn, W. J. Randomised clinical trial: a phase 1b study of GB004, an oral HIF-1alpha stabiliser, for treatment of ulcerative colitis. Aliment Pharmacol Ther 55, 401–411, doi:10.1111/apt.16753 (2022).

8 Chakraborty, A. A., Laukka, T., Myllykoski, M., Ringel, A. E., Booker, M. A., Tolstorukov, M. Y., Meng, Y. J., Meier, S. R., Jennings, R. B., Creech, A. L., Herbert, Z. T., McBrayer, S. K., Olenchock, B. A., Jaffe, J. D., Haigis, M. C., Beroukhim, R., Signoretti, S., Koivunen, P. & Kaelin, W. G., Jr. Histone demethylase KDM6A directly senses oxygen to control chromatin and cell fate. Science 363, 1217–1222, doi:10.1126/science.aaw1026 (2019).

9 Prickaerts, P., Adriaens, M. E., Beucken, T. V. D., Koch, E., Dubois, L., Dahlmans, V. E. H., Gits, C., Evelo, C. T. A., Chan-Seng-Yue, M., Wouters, B. G. & Voncken, J. W. Hypoxia increases genome-wide bivalent epigenetic marking by specific gain of H3K27me3. Epigenetics Chromatin 9, 46, doi:10.1186/s13072-016-0086-0 (2016).

10 Caldwell, C. C., Kojima, H., Lukashev, D., Armstrong, J., Farber, M., Apasov, S. G. & Sitkovsky, M. V. Differential effects of physiologically relevant hypoxic conditions on T lymphocyte development and effector functions. J Immunol 167, 6140–6149, doi:10.4049/jimmunol.167.11.6140 (2001).

11 Lukashev, D., Klebanov, B., Kojima, H., Grinberg, A., Ohta, A., Berenfeld, L., Wenger, R. H., Ohta, A. & Sitkovsky, M. Cutting edge: hypoxia-inducible factor 1alpha and its activation-inducible short isoform I.1 negatively regulate functions of CD4+ and CD8+ T lymphocytes. J Immunol 177, 4962–4965, doi:10.4049/jimmunol.177.8.4962 (2006).

12 Shehade, H., Acolty, V., Moser, M. & Oldenhove, G. Cutting Edge: Hypoxia-Inducible Factor 1 Negatively Regulates Th1 Function. J Immunol 195, 1372–1376, doi:10.4049/jimmunol.1402552 (2015).

13 Westendorf, A. M., Skibbe, K., Adamczyk, A., Buer, J., Geffers, R., Hansen, W., Pastille, E. & Jendrossek, V. Hypoxia Enhances Immunosuppression by Inhibiting CD4+ Effector T Cell Function and Promoting Treg Activity. Cell Physiol Biochem 41, 1271–1284, doi:10.1159/000464429 (2017).

14 Powrie, F. T cells in inflammatory bowel disease: protective and pathogenic roles. Immunity 3, 171–174, doi:10.1016/1074-7613(95)90086-1 (1995).

15 Powrie, F., Leach, M. W., Mauze, S., Menon, S., Caddle, L. B. & Coffman, R. L. Inhibition of Th1 responses prevents inflammatory bowel disease in scid mice reconstituted with CD45RBhi CD4+ T cells. Immunity 1, 553–562, doi:10.1016/1074-7613(94)90045-0 (1994).

16 Sartor, R. B. Mechanisms of disease: pathogenesis of Crohn’s disease and ulcerative colitis. Nat Clin Pract Gastroenterol Hepatol 3, 390–407, doi:10.1038/ncpgasthep0528 (2006).

17 Neildez-Nguyen, T. M. A., Bigot, J., Da Rocha, S., Corre, G., Boisgerault, F., Paldi, A. & Galy, A. Hypoxic culture conditions enhance the generation of regulatory T cells. Immunology 144, 431–443, doi:10.1111/imm.12388 (2015).

18 Roman, J., Rangasamy, T., Guo, J., Sugunan, S., Meednu, N., Packirisamy, G., Shimoda, L. A., Golding, A., Semenza, G. & Georas, S. N. T-cell activation under hypoxic conditions enhances IFN-gamma secretion. Am J Respir Cell Mol Biol 42, 123–128, doi:10.1165/rcmb.2008-0139OC (2010).

19 Rutz, S., Kayagaki, N., Phung, Q. T., Eidenschenk, C., Noubade, R., Wang, X., Lesch, J., Lu, R., Newton, K., Huang, O. W., Cochran, A. G., Vasser, M., Fauber, B. P., DeVoss, J., Webster, J., Diehl, L., Modrusan, Z., Kirkpatrick, D. S., Lill, J. R., Ouyang, W. & Dixit, V. M. Deubiquitinase DUBA is a post-translational brake on interleukin-17 production in T cells. Nature 518, 417–421, doi:10.1038/nature13979 (2015).

20 Kruidenier, L., Chung, C. W., Cheng, Z., Liddle, J., Che, K., Joberty, G., Bantscheff, M., Bountra, C., Bridges, A., Diallo, H., Eberhard, D., Hutchinson, S., Jones, E., Katso, R., Leveridge, M., Mander, P. K., Mosley, J., Ramirez-Molina, C., Rowland, P., Schofield, C. J., Sheppard, R. J., Smith, J. E., Swales, C., Tanner, R., Thomas, P., Tumber, A., Drewes, G., Oppermann, U., Patel, D. J., Lee, K. & Wilson, D. M. A selective jumonji H3K27 demethylase inhibitor modulates the proinflammatory macrophage response. Nature 488, 404–408, doi:10.1038/nature11262 (2012).

21 Wei, G., Wei, L., Zhu, J., Zang, C., Hu-Li, J., Yao, Z., Cui, K., Kanno, Y., Roh, T. Y., Watford, W. T., Schones, D. E., Peng, W., Sun, H. W., Paul, W. E., O’Shea, J. J. & Zhao, K. Global mapping of H3K4me3 and H3K27me3 reveals specificity and plasticity in lineage fate determination of differentiating CD4+ T cells. Immunity 30, 155–167, doi:10.1016/j.immuni.2008.12.009 (2009).

22 Wei, J., Duramad, O., Perng, O. A., Reiner, S. L., Liu, Y. J. & Qin, F. X. Antagonistic nature of T helper 1/2 developmental programs in opposing peripheral induction of Foxp3+ regulatory T cells. Proc Natl Acad Sci U S A 104, 18169–18174, doi:10.1073/pnas.0703642104 (2007).

23 Bradley, D., Smith, A. J., Blaszczak, A., Shantaram, D., Bergin, S. M., Jalilvand, A., Wright, V., Wyne, K. L., Dewal, R. S., Baer, L. A., Wright, K. R., Stanford, K. I., Needleman, B., Brethauer, S., Noria, S., Renton, D., Joseph, J. J., Lovett-Racke, A., Liu, J. & Hsueh, W. A. Interferon gamma mediates the reduction of adipose tissue regulatory T cells in human obesity. Nat Commun 13, 5606, doi:10.1038/s41467-022-33067-5 (2022).

24 Mottet, C., Uhlig, H. H. & Powrie, F. Cutting edge: cure of colitis by CD4+CD25+ regulatory T cells. J Immunol 170, 3939–3943, doi:10.4049/jimmunol.170.8.3939 (2003).

25 Mayne, C. G. & Williams, C. B. Induced and natural regulatory T cells in the development of inflammatory bowel disease. Inflamm Bowel Dis 19, 1772–1788, doi:10.1097/MIB.0b013e318281f5a3 (2013).

26 Yamada, A., Arakaki, R., Saito, M., Tsunematsu, T., Kudo, Y. & Ishimaru, N. Role of regulatory T cell in the pathogenesis of inflammatory bowel disease. World J Gastroenterol 22, 2195–2205, doi:10.3748/wjg.v22.i7.2195 (2016).

27 Powrie, F., Carlino, J., Leach, M. W., Mauze, S. & Coffman, R. L. A critical role for transforming growth factor-beta but not interleukin 4 in the suppression of T helper type 1-mediated colitis by CD45RB(low) CD4+ T cells. J Exp Med 183, 2669–2674, doi:10.1084/jem.183.6.2669 (1996).

28 Gozdecka, M., Meduri, E., Mazan, M., Tzelepis, K., Dudek, M., Knights, A. J., Pardo, M., Yu, L., Choudhary, J. S., Metzakopian, E., Iyer, V., Yun, H., Park, N., Varela, I., Bautista, R., Collord, G., Dovey, O., Garyfallos, D. A., De Braekeleer, E., Kondo, S., Cooper, J., Gottgens, B., Bullinger, L., Northcott, P. A., Adams, D., Vassiliou, G. S. & Huntly, B. J. P. UTX-mediated enhancer and chromatin remodeling suppresses myeloid leukemogenesis through noncatalytic inverse regulation of ETS and GATA programs. Nat Genet 50, 883–894, doi:10.1038/s41588-018-0114-z (2018).

29 Wang, C., Lee, J. E., Cho, Y. W., Xiao, Y., Jin, Q., Liu, C. & Ge, K. UTX regulates mesoderm differentiation of embryonic stem cells independent of H3K27 demethylase activity. Proc Natl Acad Sci U S A 109, 15324–15329, doi:10.1073/pnas.1204166109 (2012).

30 Wang, S. P., Tang, Z., Chen, C. W., Shimada, M., Koche, R. P., Wang, L. H., Nakadai, T., Chramiec, A., Krivtsov, A. V., Armstrong, S. A. & Roeder, R. G. A UTX-MLL4-p300 Transcriptional Regulatory Network Coordinately Shapes Active Enhancer Landscapes for Eliciting Transcription. Mol Cell 67, 308–321 e306, doi:10.1016/j.molcel.2017.06.028 (2017).

31 Ke, X., Chen, C., Song, Y., Cai, Q., Li, J., Tang, Y., Han, X., Qu, W., Chen, A., Wang, H., Xu, G. & Liu, D. Hypoxia modifies the polarization of macrophages and their inflammatory microenvironment, and inhibits malignant behavior in cancer cells. Oncol Lett 18, 5871–5878, doi:10.3892/ol.2019.10956 (2019).

32 Peng, W., Xie, Y., Luo, Z., Liu, Y., Xu, J., Li, C., Qin, T., Lu, H. & Hu, J. UTX deletion promotes M2 macrophage polarization by epigenetically regulating endothelial cell-macrophage crosstalk after spinal cord injury. J Nanobiotechnology 21, 225, doi:10.1186/s12951-023-01986-0 (2023).

33 Breese, E., Braegger, C. P., Corrigan, C. J., Walker-Smith, J. A. & MacDonald, T. T. Interleukin-2- and interferon-gamma-secreting T cells in normal and diseased human intestinal mucosa. Immunology 78, 127–131 (1993).

34 Fujino, S., Andoh, A., Bamba, S., Ogawa, A., Hata, K., Araki, Y., Bamba, T. & Fujiyama, Y. Increased expression of interleukin 17 in inflammatory bowel disease. Gut 52, 65–70, doi:10.1136/gut.52.1.65 (2003).

35 Ito, H. & Fathman, C. G. CD45RBhigh CD4+ T cells from IFN-gamma knockout mice do not induce wasting disease. J Autoimmun 10, 455–459, doi:10.1016/s0896-8411(97)90152-9 (1997).

36 Liu, Z. J., Yadav, P. K., Su, J. L., Wang, J. S. & Fei, K. Potential role of Th17 cells in the pathogenesis of inflammatory bowel disease. World J Gastroenterol 15, 5784–5788, doi:10.3748/wjg.15.5784 (2009).

37 Sin, J. H., Zuckerman, C., Cortez, J. T., Eckalbar, W. L., Erle, D. J., Anderson, M. S. & Waterfield, M. R. The epigenetic regulator ATF7ip inhibits Il2 expression, regulating Th17 responses. J Exp Med 216, 2024–2037, doi:10.1084/jem.20182316 (2019).

38 O’Connor, W., Jr., Kamanaka, M., Booth, C. J., Town, T., Nakae, S., Iwakura, Y., Kolls, J. K. & Flavell, R. A. A protective function for interleukin 17A in T cell-mediated intestinal inflammation. Nat Immunol 10, 603–609, doi:10.1038/ni.1736 (2009).

39 Ogawa, A., Andoh, A., Araki, Y., Bamba, T. & Fujiyama, Y. Neutralization of interleukin-17 aggravates dextran sulfate sodium-induced colitis in mice. Clin Immunol 110, 55–62, doi:10.1016/j.clim.2003.09.013 (2004).

40 Ikejiri, A., Nagai, S., Goda, N., Kurebayashi, Y., Osada-Oka, M., Takubo, K., Suda, T. & Koyasu, S. Dynamic regulation of Th17 differentiation by oxygen concentrations. Int Immunol 24, 137–146, doi:10.1093/intimm/dxr111 (2012).

41 Ostanin, D. V., Bao, J., Koboziev, I., Gray, L., Robinson-Jackson, S. A., Kosloski-Davidson, M., Price, V. H. & Grisham, M. B. T cell transfer model of chronic colitis: concepts, considerations, and tricks of the trade. Am J Physiol Gastrointest Liver Physiol 296, G135–146, doi:10.1152/ajpgi.90462.2008 (2009).

42 Cook, K. D., Shpargel, K. B., Starmer, J., Whitfield-Larry, F., Conley, B., Allard, D. E., Rager, J. E., Fry, R. C., Davenport, M. L., Magnuson, T., Whitmire, J. K. & Su, M. A. T Follicular Helper Cell-Dependent Clearance of a Persistent Virus Infection Requires T Cell Expression of the Histone Demethylase UTX. Immunity 43, 703–714, doi:10.1016/j.immuni.2015.09.002 (2015).

43 Read, S. & Powrie, F. Induction of inflammatory bowel disease in immunodeficient mice by depletion of regulatory T cells. Curr Protoc Immunol Chapter 15, Unit 15 13, doi:10.1002/0471142735.im1513s30 (2001).

